# Determination of the Optical Properties of Thylakoid Membranes from Whole Leaf Reflectance Measurements Using a Capacitor and Unbound Electron Model

**DOI:** 10.1101/2021.06.30.450215

**Authors:** Ranjan S. Muttiah, Sergey N. Savenkov

## Abstract

This paper demonstrates that a capacitor equivalent along with unbound electrons can be used to model thylakoid membranes in grana stacks. From whole leaf reflectance measurements at normal incidences at 660nm wavelength and taken from the literature (Hu et al. 2020), refractive indices are obtained from the Fresnel’s equation for transverse electric (TE) and transverse magnetic (TM) polarization. The TE and TM polarizations for external reflectance depict the Brewster angle at which the magnitude of the reflected electric vector is zero; the internal reflections show that there’s a narrow angle window of about 10 degrees before the internally refracted light goes into critical angle. The clustering and separation of reflection measurements with angle of incidence is explained using Fresnel’s equation; the cross-over angle beyond the Brewster’s angle for internal reflection at which reflectance measurements separate with incidence angle is given by 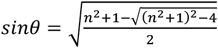. The predicted relaxation times from a capacitor and unbound electron model gave favorable comparisons against commonly reported fluorescence times in the 0.1 to 1 ns range (our results gave 0.5-0.8 ns). The di-electric constant for the membrane is estimated to be 5. The stacking number (number of grana layers) is consistent with the light penetration depth (skin depth). The magnetic permeability was shown to be close to that of vacuum and therefore the thylakoid lacks any magnetic properties as would be expected for such a transparent media. An *in-vivo* estimate based on thermal equilibrium of molecules for the permanent dipole moment of the chlorophyll molecule gave 2,025D (Debye).

## Introduction

The structure of photosynthetic apparatuses and light capture events during photosynthesis has had an exciting and enriching history (Seliger and McElroy, 1965;Govindjee et al., 2005; Nobel, 2020). In this paper, we use one of the most profound of those discoveries, namely Maxwell’s equations, to derive certain electrical parameters for the thylakoid membrane relying on measurements and inferences made from across a variety of observation scales ranging from the leaf level to the quantum level. Due to the complexity of this topic, we have used observations from different sources to determine the input to equations derived from Maxwell’s equations i.e. a single specimen subject to observation at all scales is simply not available (perhaps even impossible) and a fundamental assumption here is that measurements made from different specimens and plant species generally holds across photosynthesis of all higher plants with leaves.

### Theory & Background

An optical parameter that connects the macro world to the micro-world is the refractive index. From Maxwell’s equations and continuity conditions at the boundary between two interfaces, the following well-known Fresnel equations between angle of incidence and amplitudes of reflected and incident assuming Snell’s law is obtained for the transverse electric (TE) and magnetic polarizations (TM) (Fowles, 1975):

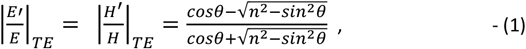

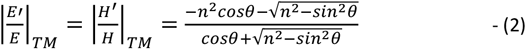

Where, E and H are the magnitudes of the electric and magnetic vectors, the prime notation is for the reflected electric and magnetic vectors (double ‘‘ prime is for refracted vectors), n is the refractive index (n=n_2_/n_1_) given as the ratio of refractive indices between the refracting media (n_2_) and incident media (n_1_), and θ is the angle of incidence. When the angle of incidence is zero, the index of refraction is be obtained from:

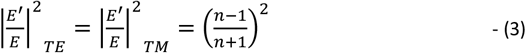

These ratios are the ratios of the light intensities that can be measured from reflectance measurements on whole leaves to obtain the refractive index; we used it to determine the refractive index at the red chlorophyll absorption peak at 660 nm from Hu et al. (2020). When 1/n is used instead of n in equations (1) and (2) the reflectance is called internal reflectance and signifies the light refraction and reflection from a the media with higher refraction index to that of the lower refractive index i.e., a light beam shone within the leaf to the outside. When the external and internal reflections are in each polarizations are equal, there is a cross-over angle prior to internally reflected light from within the media going into critical angle (see Appendix A for the derivation). The reflection measurements made beyond the pigment absorbing near-infra red ranges were assumed to be due to the influence of water, although other leaf internal structures also contribute to reflectance in the near infra-red range (see Hunt and Rock, 1989) and a refractive index of 1.33 was used for water.

After having obtained the refractive index at 600nm from whole leaf measurements, the next step was to use the relevant equations from EM theory to model the electron transport process across the thylakoid membranes in leaves, subject to assumptions on light capture and “free” electron release from one side of the thylakoid membrane to another as being represented by fluorescence emission following light capture (after the e-transport process was complete). Within the chloroplasts of plant leaves, the thylakoid membranes are packed into grana stacks; the grana stacks are connected to each other through stroma lamellae (Nobel, 2020).

Chlorophyll molecules which absorb in the blue and red parts of the visible spectrum transmit the captured photon energy via resonance transfer to PSII molecules. At the PSII molecule (which is activated at 680 nm), the electron is transported through a sequence of acceptor-donor exchanges before the e-reaches the PSI molecule (which is activated by 700nm). In this paper, the light capture and electron transfer across the thylakoid membrane of a single grana disc was modeled as a capacitor with a dielectric substance as conceptualized in Figure 1. This represented the initial phase (before action phase when chemical reactions take place) during which light is captured, energy is transferred by resonance from chlorophyll molecules to PSII in a light harvesting configuration, and ultimately the e-are transferred from the inner side of the thylakoid membrane to the outer side where chemical reactions (ATP generation, NADPH reduction etc.) take place. The end of the e-transfer process is signified by fluorescence decay (ns range) after which the chemical reactions take place over a much longer time period (ms range). It should be noted that not all the energy captured by the chlorophyll molecules activate the electrons to excited singlet spin states; much of the energy is also accounted for by the interaction between electrons and the nuclei as described by the Frank-Condon progression (Nobel, 2020; Quantum Chemistry, LibreTexts, 2021).

**Figure 1.**
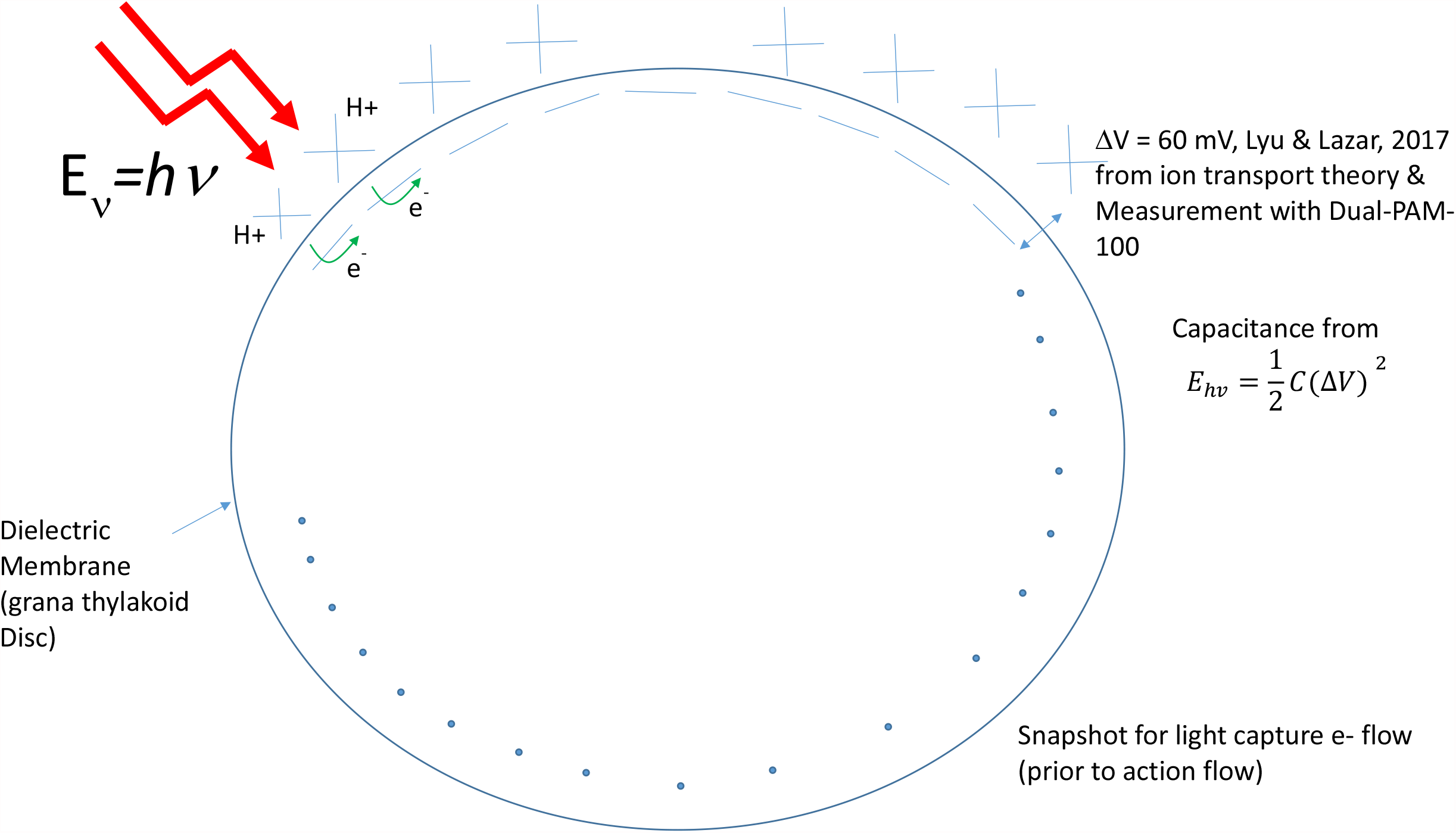
Capacitor model for a thylakoid disc modeled as a sphere-equivalent dielectric membrane.

Lyu and Lazar (2017) performed electrochromic shift (ECS) measurements using a Dual-PAM-100 system at two different wavelengths at 515 and 535 nm to capture the change in transmittance across those wavelengths and estimated the associated potential difference across the thylakoid membrane from the timing difference by fitting an ionic transport model. They found a peak potential difference of 60 mV and this was used in this paper.

The velocity (**v**) of unbound conduction electrons in the midst of the light’s electric field (**E**) are modeled through the following differential equation (Fowles, 1975; the steps here follow those of Fowles and are reproduced for convenience of reader):

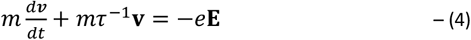

Where, m is the mass of an electron and τ is the relaxation time which together (mτ^-1^) represent a frictional dissipation constant. Current density J is given by:

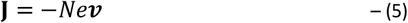

Where, N is the number of conduction electrons per unit volume. Substituting equation (5) in (4) gives:

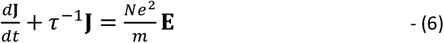

The homogeneous form of this equation is when the left hand side is zero giving the transient current decay as:

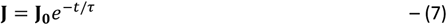

Thus, from this equation it is seen that τ represents the exponential decay in the electrical current from some initial value. Fluorescence decay is given by a linear rate constant equation for electrons activated in their π-orbitals given by (Nobel, 2020):

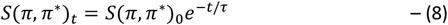

It is understood that while equation (6) describes unbound and free electron moving in an electric field, and electron motion in the thylakoid membrane involves the transfer of electron from one molecule to another through an acceptor-donor process, it is assumed that effectively the *two processes are equivalent* from a modeling perspective.

For a static electric field, equation (6) takes the form:

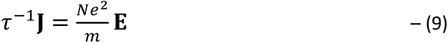

Comparing equation (9) to the definition of current density **J** = *σ***E** gives the static conductivity σ as:

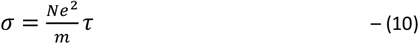

Assuming a harmonic wave solution for both the electric field and current density, and solving for current density J gives:

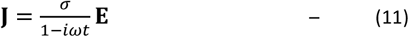

The Maxwell’s wave equations in terms of the electric vector and current density is given by the following expression after ignoring polarization effects from bound electrons:

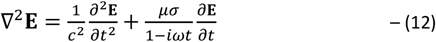

Where μ_0_ is the permeability of the media (thylakoid membrane). A plane harmonic wave of the form *E* = *E*_0_*e*^*i*(*ξz*−*ω t*)^ is now tried for solution of above equation assuming no split in the wave vector due to anisotropy (isotropic conditions at the boundary), where ξ is a wave vector represented by complex number for the wave vector (k) and extinction coefficient α through:

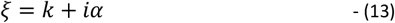

By use of the operator 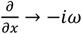, the following relationship is obtained:

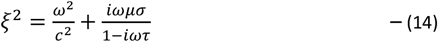

At low frequencies, equation (14) can be approximated by:

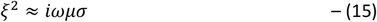

Substituting the complex number from (13) for ξ and equating the real part to zero and imagery part to the RHS, the above equation gives:

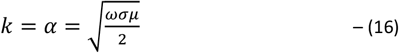

The complex wave number ξ is related to the complex refractive index η (= *n* + *i χ*) via:

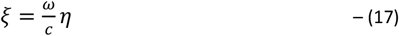

Equating the real parts to each other, and the imagery parts to each other gives:

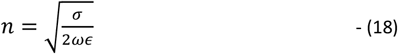

Where, ε is the permittivity of the media. The amplitude of the electric vector drops by an exponential factor from the surface (“skin depth”) according to the following:

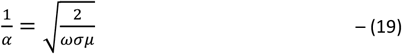

The critical parameter required to obtain the relaxation time τ is the number of charges per unit volume N in units of charges/m^3^. This was obtained by modeling the thylakoid membrane as a di-electric material with charges on the inside and outside of the membranes, representing the vector transport of electrons from the side of the thylakoid in lumen to the outside of the thylakoid membrane immersed in stroma (see Figure 5-18, Nobel, 2020). As in Nobel (2020) we used a capacitor to model the di-electric using the equation for the energy content from photon absorption via the relationship:

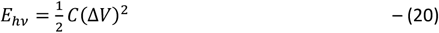

Where, C is the capacitance and ΔV is the voltage difference across the thylakoid membrane which was taken from Lyu and Lazar (2017). To get the charge per unit volume, a grana disc with thylakoid membranes was transformed from a cylindrical shape to a sphere equivalent shape via the volume and area equations:

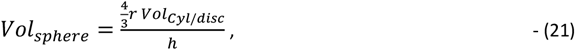

And the surface area:

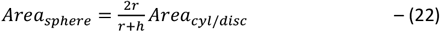

The dimensions for the grana disc were obtained from Austin and Staehlin (2011). It was also assumed that there were 250 chlorophyll molecules in a light harvesting complex (Malkin and Niyogi, 2000) with each chlorophyll transferring the captured photon energy via resonance transfer to a PSII molecule.

The thylakoid permittivity was calculated from the capacitance equation:

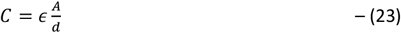

Where, A is the area of charge and d is the charge separation distance (taken as the thickness of the thylakoid membrane). The magnetic permeability was calculated via the equation (Fowles, 1975):

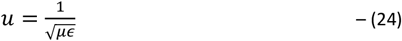

Where u is the speed of light within the media which was obtained from the refractive index via:

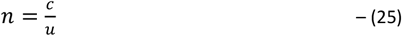

Where, c is the speed of light (3.0×10^8^ m/s).

## Material & Methods

Leaf level reflectance measurements made by Hu et al., (2020) on five different types of leaves from 4 species found in Beijing was used in this paper; reflectance at normal incidence at 660 for the strong absorption band of chlorophyll molecules are shown in Table 1. The experimental set up consisted of a tunable and collimated laser operating at 2 Watt transmission power, mirror collector, and photodiode detector which was connected to an oscilloscope for data recording (the reader is referred to their paper for additional experimental details). The refractive indices on application of equation (3) are shown in Table 1. These values are comparable to the refractive indices obtained from chlorophyll using a bound electron model by Achi and Castro (2021). Microscopic images of grana stack from Austin and Staehlin (2011) was used to determine the average height of a grana disc to be 0.118 μm, and the radius of a grana disc to be 0.75 μm. The height was obtained by counting the number of grana layers and then dividing by the total height measured from the image using the provided scale bar.

**Table 1.** Reflectance at normal incidence from Hu et al. (2020). All values shown are for normal incidences. [Double check these values!!].

| Sample | Family Name | 660 nm reflectance | n | Brewster angle, deg | Cross over angle, deg |
| --- | --- | --- | --- | --- | --- |
| Fraxinus pennsylvanica (green leaf) | Olive | 0.145 | 2.23 | 65.8 | 23.8 |
| Fraxinus pennsylvanica (yellow leaf) | Olive | 0.08 | 1.79 | 60.8 | 28.8 |
| Eucommia ulmoides | Eucommiaceae | 0.11 | 2.0 | 63.4 | 26.2 |
| Amygdalus triloba | Rose | 0.08 | 1.79 | 60.8 | 28.8 |
| Magnolia denudate | Magnoliaceae | 0.075 | 1.76 | 60.4 | 29.2 |

## Results & Discussion

Figure 2 give the results of applying Fresnel’s equation for the 660 nm wavelength whole leaf measurement made on the Fraxinus green leaf sample (other samples gave similar plots and can be found in the supplemental material) and for water; the associated Brewster angle at which the external and internal reflection TM magnitudes are 0 given in Table 1; the cross-over angle (Table 1) and the Brewster angle are smaller for the membrane (the 660nm light on leaf at normal incidence was assumed to reach the thylakoid membrane) than for water. It’s also interesting to note that the water critical angle is very near the Brewster angle for external reflection. These graphs can be transformed to capture the polarization changes on the viewing side (air viewing side) by accounting for the wave vectors (k, k’, k’’) and assuming a light source within the leaf to simulate reflected light from within the leaf (see calculations in the supplemental material spreadsheet). The plot so obtained is shown in Figure 3. We use these plots to explain the observed clustering before the cross-over angles and beyond the cross-over angles as found in the Hu et al. (2020) plots where reflectance measurements up to a certain incident angle cluster together and after which the reflectance measurements start diverging. We explain this as due to enhancement of polarizations prior to the cross-over angles as shown in Figure 2b.

**Figure 2.**
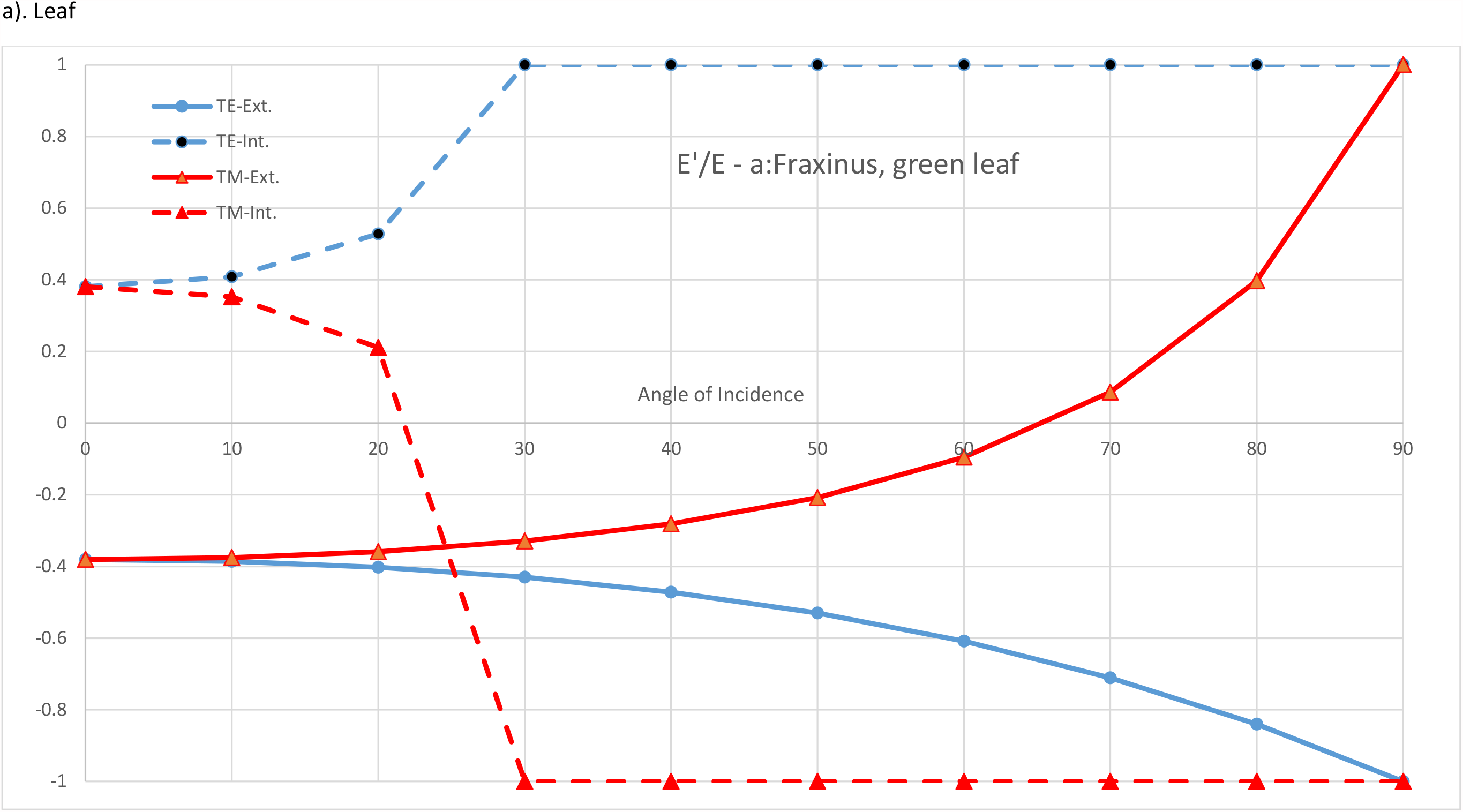
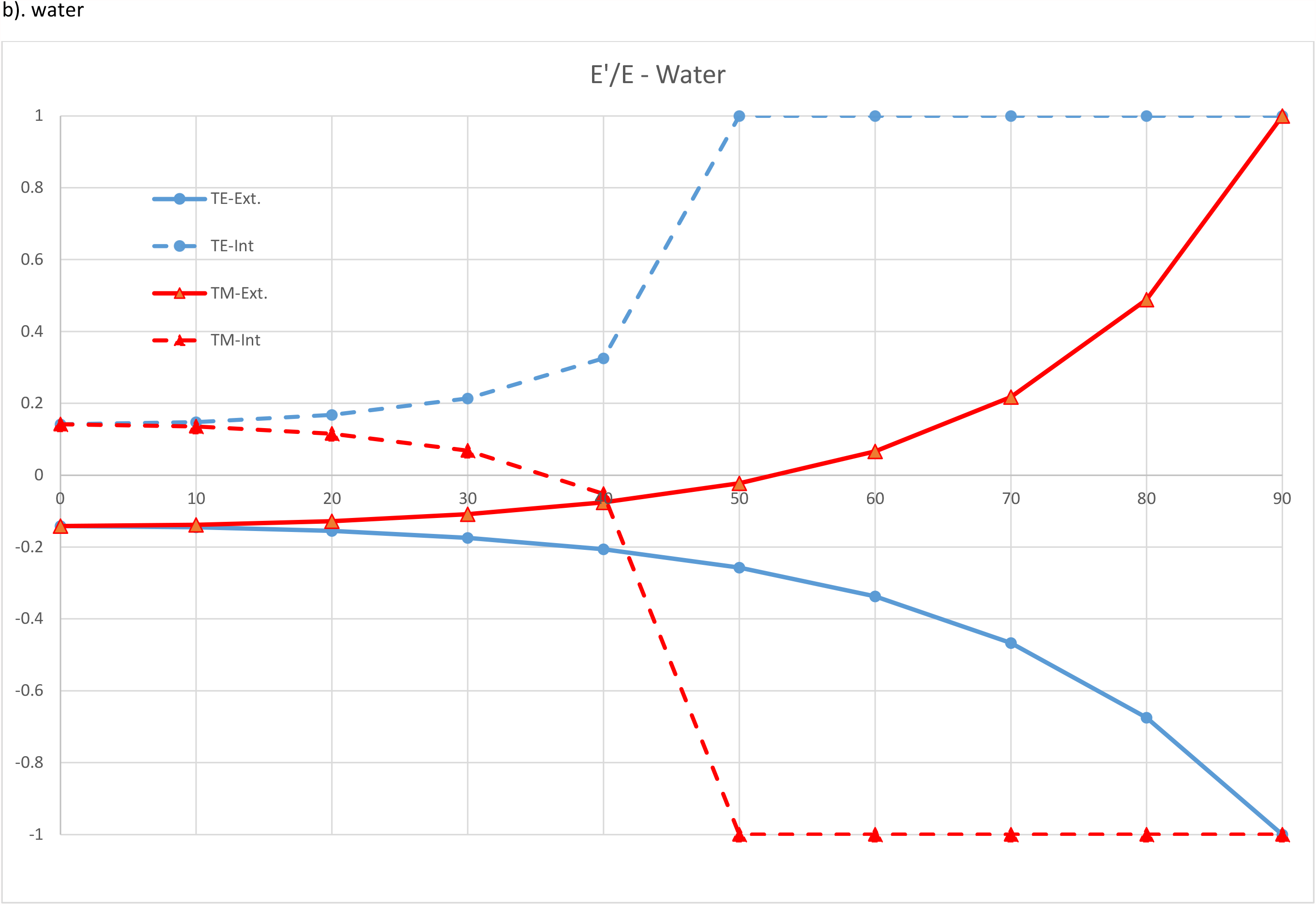
Reflected to transmitted electric field for external reflectance n=n_2_/n_1_ > 1 and internal n < 1 reflectance at 660nm for a). leaf and b). water.

**Figure 3.**
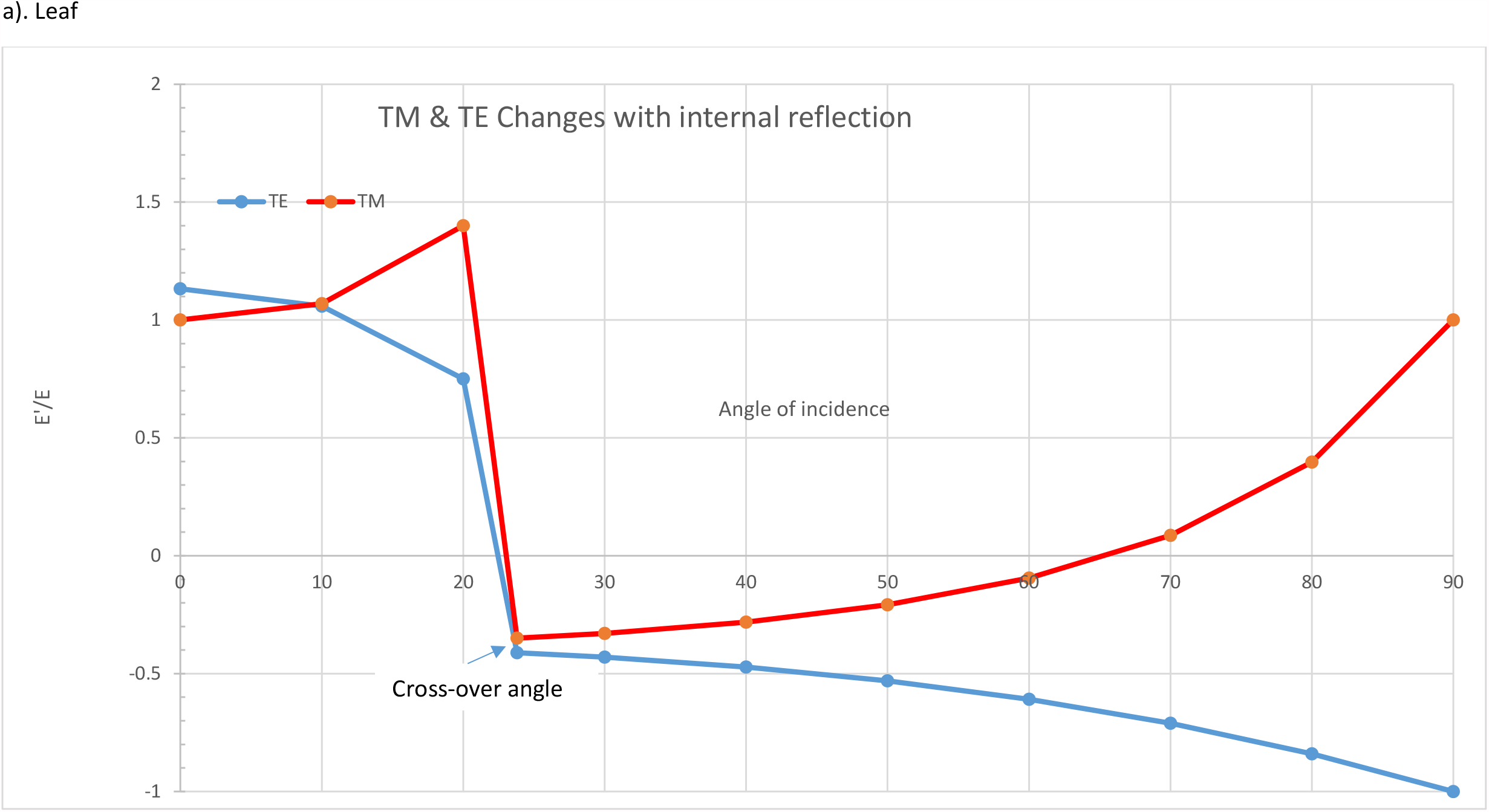
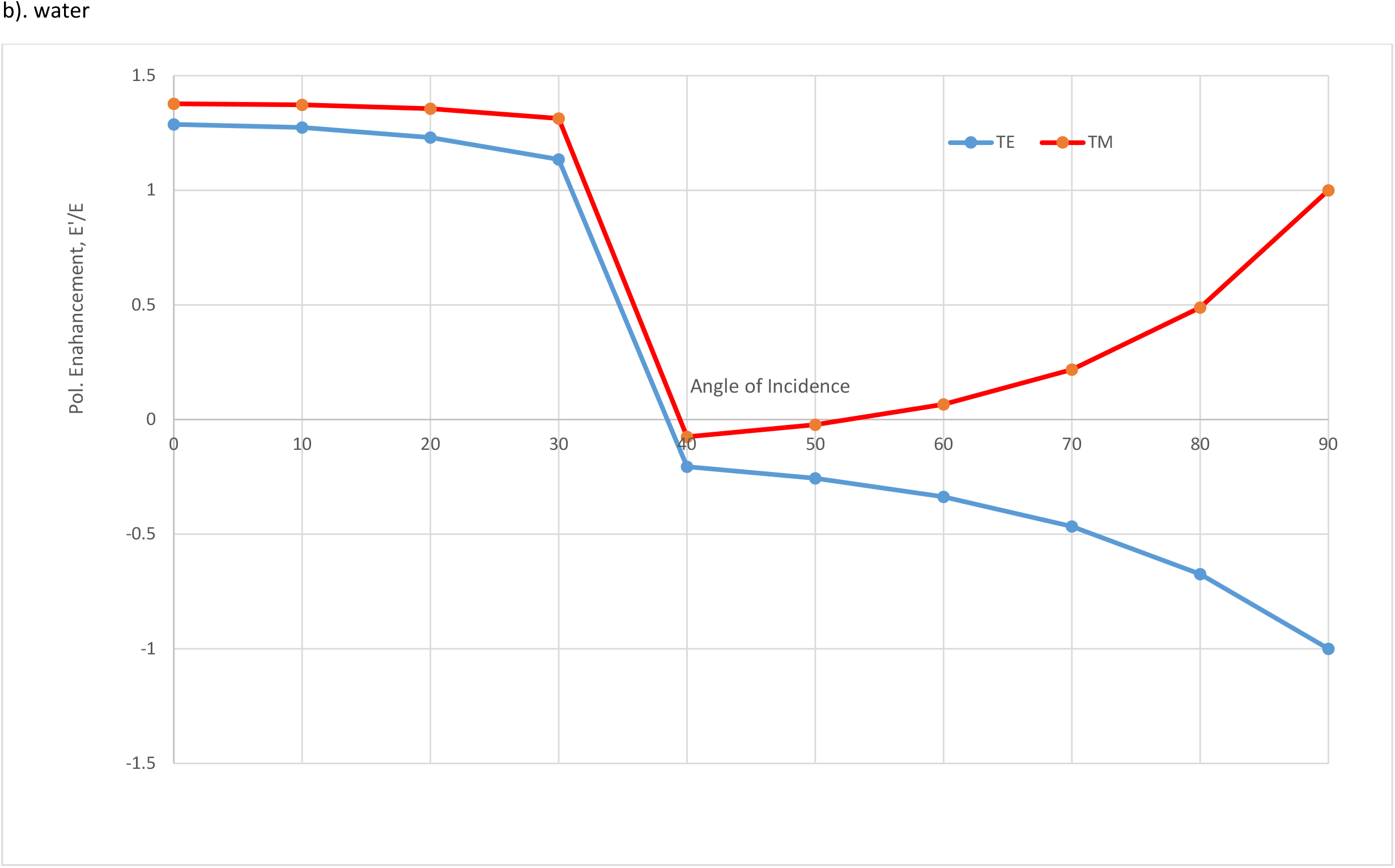
Polarization enhancement (magnitude of TE and TM polarization with internal reflection)

Results from the capacitance calculations are stepped through below; the reader can examine these calculations in the supplemental material spreadsheet under the tab “Capacitor Model.”

1. The energy per mole of red light photons at 660nm was:

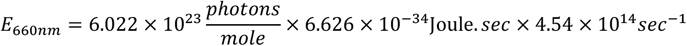

Giving:

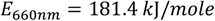
2. Assuming 250 chlorophyll molecules per light harvesting complex:

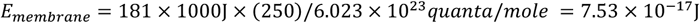
3. The voltage difference across the thylakoid membrane during the electron transfer process from the inner lumen facing to the outer stroma facing thylakoid membrane was taken as 60mV as given in Lyu and Lazar (2017).
4. Substituting terms in equation (19) gives the capacitance as: 4.183 × 10^−14^Farad.
5. The equivalent area of a thylakoid sphere based on measurements made from Austin and Staehlin (2011) in equation (21) gave: 7.07 × 10^−12^m^2^.
6. Dividing step 4 and 5 gives: 5.92 mF/m^2^.
7. The charge concentration c’ in charges/mole inside sphere:

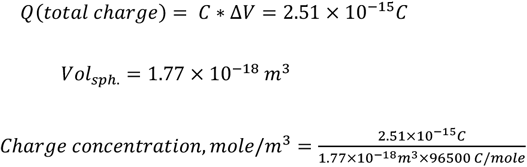

Where, 96,500 C/mole is the Faraday constant.

Step 8). The number of charges is obtained by multiplying the above number by the Avagadro’s number giving N:

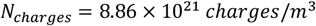

Step 9). The dielectric constant and from it the permittivity and permeability were obtained as follows.

Thylakoid membrane thickness was taken at 7.5nm (Nobel, 2020).

Substituting for values in Eqn (23) gave:

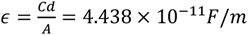

Dividing by the vacuum permittivity gives the di-electric constant as 5.015.

Using as example a refractive index of 2.23 for Fraxinus green leaf in Eqns(24) and (25) gives, the magnetic permeability as:

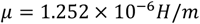

This is very near the vacuum permittivity of 1.2566 × 10^−6^ H/m demonstrating the non-magnetic nature of the thylakoid media. Therefore, the di-electric constant can also be calculated directly from the refractive index via (Fowles, 1975): 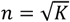. Substituting a 2.23 gives K as 4.97 which is close that obtained from the capacitance equation.

Step 10). The next step involved substituting N in equation (10), current density from (10) into equation (17) and adjusting the relaxation time τ until the refractive index n matched the values in Table 1. These results are given in Table 2 for the five leaf samples. The relaxation times obtained fall within those reported by Nobel (2020) of 0.15ns to Malkin and Niyogi who states in the ns range (presumably within 0.1-1 ns).

**Table 2.** Relaxation times, current density and skin depth obtained from the steps given in the results section.

| Sample | Family Name | Relaxation time, ns | Current density, $\sigma$ mho/m | Skin depth, $\mu\text{m}$ | Predicted Layers |
| --- | --- | --- | --- | --- | --- |
| Fraxinus pennsylvanica (green leaf) | Olive | 0.8 | $2.02 \times 10^5$ | 0.13 | 18 |
| Fraxinus pennsylvanica (yellow leaf) | Olive | 0.6 | $1.513 \times 10^5$ | 0.15 | 20 |
| Eucommia ulmoides | Eucommiaceae | 0.71 | $1.8 \times 10^5$ | 0.14 | 19 |
| Amygdalus triloba | Rose | 0.52 | $1.31 \times 10^5$ | 0.16 | 22 |
| Magnolia denudate | Magnoliaceae | 0.6 | $2.42 \times 10^4$ | 0.15 | 20 |

Step 11). The skin depth was obtained Eqn (19) and the number of predicted optically active thylakoid layers was obtained by dividing the skin depth by the thickness of thylakoid membrane. These results are shown in Table 2 (reader is referred to the supplementary spreadsheet). Austin and Staehlin (2011) in their detailed 3D imaging of grana found that a stacking number of 17. Our work suggests that the stacking number (Table 2) is predictable from the skin depth.

The determination of the di-electric constant enables estimating the dipole moment for the chlorophyll molecule *in vivo* via the following formula which accounts for thermal motion on dielectric properties (Feynman, 1963, Book II, formula 11.21):

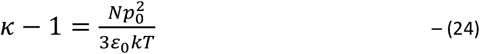

Where, k is the Boltzmann constant, T the temperature in Kelvins and p_0_ the permanent dipole. Substituting values gives p_0_ = 2,025D where D is the Debye unit (=3.34×10^−30^ C.m); water by comparison has a p_0_ of 1.85D. This demonstrates that the dipole moment in the porphyrin ring of the chlorophyll molecule has strong alignment with the electric field during light capture.

## Supporting information

Supplemental Calcs

## Appendix A

Equating equations (1) and (2) for the TE external and TM internal reflectance polarizations after substituting 1/n for refractive index for the internal reflectance and multiplying numerator and denominator by n^2^ gives:

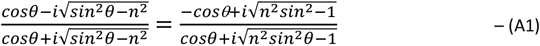

Multiplying the numerator and denominator by the denominator complex conjugates, simplifying, and then equating the real part (0 on the RHS) gives:

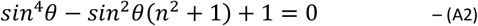

Solving for physically realizable solutions:

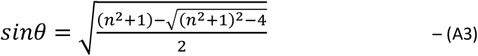

